# Developing folate-conjugated miR-34a therapeutic for prostate cancer treatment: Challenges and promises

**DOI:** 10.1101/2023.11.25.568612

**Authors:** Wen (Jess) Li, Yunfei Wang, Xiaozhuo Liu, Shan Wu, Moyi Wang, Steven G. Turowski, Joseph A. Spernyak, Amanda Tracz, Ahmed M. Abdelaal, Kasireddy Sudarshan, Igor Puzanov, Gurkamal Chatta, Andrea L. Kasinski, Dean G. Tang

## Abstract

Prostate cancer (PCa) remains a common cancer with high mortality in men due to its heterogeneity and the emergence of drug resistance. A critical factor contributing to its lethality is the presence of prostate cancer stem cells (PCSCs), which can self-renew, long-term propagate tumors and mediate treatment resistance. MicroRNA-34a (miR-34a) has shown promise as an anti-PCSC therapeutic by targeting critical molecules involved in cancer stem cell (CSC) survival and functions. Despite extensive efforts, the development of miR-34a therapeutics still faces challenges, including non-specific delivery and delivery-associated toxicity. One emerging delivery approach is ligand-mediated conjugation, aiming to achieve specific delivery of miR-34a to cancer cells, thereby enhancing efficacy while minimizing toxicity. Folate-conjugated miR-34a (folate-miR-34a) has demonstrated promising anti-tumor efficacy in breast and lung cancers by targeting folate receptor α (FOLR1). Here, we first show that miR-34a, a TP53 transcriptional target, is reduced in PCa that harbors TP53 loss or mutations and that miR-34a mimic, when transfected into PCa cells, downregulated multiple miR-34a targets and inhibited cell growth. When exploring the therapeutic potential of folate-miR-34a, we found that folate-miR-34a exhibited impressive inhibitory effects on breast, ovarian and cervical cancer cells but showed minimal effects on and targeted delivery to PCa cells due to a lack of appreciable expression of FOLR1 in PCa cells. Folate-miR-34a also did not display any apparent effect on PCa cells expressing prostate-specific membrane antigen (PMSA) despite the reported folate’s binding capability to PSMA. These results highlight challenges in specific delivery of folate-miR-34a to PCa due to lack of target (receptor) expression. Our study offers novel insights on the challenges and promises within the field and cast light on the development of ligand-conjugated miR-34a therapeutics for PCa.

## 1 Introduction

Prostate cancer (PCa) remains a formidable challenge in men due to its remarkable heterogeneity and the emergence of drug resistance, resulting in ultimately lethal castration-resistant PCa called CRPC. This malignancy, characterized by multiple distinct cancer foci and varying androgen receptor (AR) expression levels, has been at the forefront of therapeutic research for decades. The current standard-of-care therapies, including androgen receptor signaling inhibitors (AR-SIs), radiotherapy and chemotherapies, have exhibited good clinical efficacy, but offering survival benefits only measured in months in advanced PCa patients [1,2]. A critical factor contributing to ARSI resistance and therapeutic failure is the existence of prostate cancer stem cells (PCSCs), a subpopulation of cells within the tumor that possess stem cell traits [3,4]. These PCSCs can long-term self-renew, propagate tumors in vivo, and are inherently ARSI-refractory. In addition, tumor progression and therapeutic treatments may induce plasticity by reprogramming non-cancer stem cells (CSCs) into PCSCs [5,6]. Consequently, PCSCs play a pivotal role in driving drug resistance and disease progression.

MicroRNAs (miRNAs), ∼22-nucleotide (nt) non-protein coding RNAs, are important posttranscriptional regulators of gene expression. MicroRNA-34a (miR-34a) is a bona fide tumor suppressor, which is downregulated in a wide range of solid tumors and hematological malignancies [7,8]. Of significance, miR-34a functions as a potent CSC suppressor by targeting key molecules essential for the survival and activities of CSCs [5,7]. In fact, extensive studies have shown that miR-34a exhibits anti-PCSC effects by targeting invasiveness and metastasis [9-11], stemness [12,13], epigenome [14,15], and cell survival [13,15,16]. Our earlier data revealed that systemic delivery of miR-34a reduced prostate tumor burden and lung metastasis by inhibiting PCSCs via targeting CD44 [11]. This indicates miR-34a as a promising therapeutic for PCSC-enriched advanced PCa.

Despite extensive translational research in the field, the development of miR-34a therapeutics has been hampered by several challenges including delivery vehicle-associated toxicity, inadequate cellular uptake and stability, and limited specificity in targeted delivery to tumors [17,18]. Current delivery strategies for miR-34a therapeutic fall into two general categories: packaged vehicles such as liposomes and nanoparticles, and vehicle-free delivery such as ligand-conjugates [5,19]. Major challenges of packaged vehicles include immunogenic effects and toxicities due to off-target effects associated with non-specific delivery [20]. To overcome these barriers, ligand-conjugation approach has been explored to achieve specific delivery of miR-34a to cancer cells, thereby enhancing therapeutic efficacy while minimizing toxicity. The concept is to directly conjugate a targeting ligand to miR-34a without a delivery vehicle. Typically, these targeting ligands are small molecules that exhibit both high affinity and specificity for receptors. On the other hand, the target receptors should be overexpressed on the surface of cancer cells relative to normal cells, and their expression level should be sufficient to enable delivery of therapeutic quantities to tumor cells. A successful example is developing folate-conjugated miR-34a (folate-miR-34a) to target breast and lung cancers via folate receptor α (FOLR1) [21]. Folate is an essential vitamin and a high-affinity ligand for the FOLR1, which is highly upregulated in ovarian, lung, breast, and other cancers [22]. Orellana et al. were the first to design and synthesize folate-miR-34a, and showed that folate-miR-34a was selectively targeted to FOLR1-expressing tumors, downregulated target genes, and suppressed the tumor growth *in vivo* in lung and breast cancers [21]. Interestingly, folate can also bind to another membrane protein prostate specific membrane antigen (PSMA), which is a clinically validated therapeutic target for PCa. PSMA is highly upregulated in PCa, and its expression has been associated with PCa progression [23-25]. ^177^Lu-PSMA-617 (Pluvicto™), which combines a PSMA-specific peptidomimetic with a therapeutical radionuclide, was the first FDA-approved PSMA-targeting therapy for metastatic PCa patients in 2022. Currently, there are several PSMA-targeting therapies undergoing clinical development, which include antibody-drug conjugates, PSMA-targeting immunotherapies, radioligand therapy, and photodynamic therapy [25,26]. With these considerations, *we hypothesized* that folate-miR-34a can be a potential therapeutic for treating PCa by targeting PCa cells expressing FOLR1 and/or PMSA.

In this study, we first show that miR-34a, a p53 transcriptional target, is significantly downregulated in PCa that have sustained p53 loss or mutations. We then show that folate-miR-34a, unexpectedly, did not elicit PCa-inhibitory effects, even in PSMA-expressing PCa cells. Further studies revealed that FOLR1, the major receptor for folate, is barely expressed in PCa. While our findings expose challenges associated with achieving specific delivery of folate-miR-34a to PCa, we provide evidence that folate-miR-34a may be a therapeutic agent for FOLR1-expressing cancers including ovarian and cervical cancers. Importantly, the insights obtained from the current study shed lights on the future development of ligand-conjugated (and unconjugated) miR-34a as potential therapeutics for advanced and aggressive PCa.

## 2 Results

### 2.1 miR-34a expression is downregulated in PCa that has TP53 loss or mutations

miR-34a is known to be a p53-regulated miRNA and a crucial component of the p53 tumor suppressor network [27-29]. Previously, we provided preliminary data showing that the expression levels of both mature miR-34a and pre-miR-34a (Figure 1A) are significantly reduced in *TP53* mutated compared to *TP53* WT prostate tumors [5]. This suggests a reciprocal association between *TP53* status and miR-34a levels in PCa. Herein, we further distilled *TP53* genetic alterations utilizing the *TCGA* (Cancer Genome Atlas) database and found that approximately 46% of primary PCa patients exhibit *TP53* alterations, including loss of heterozygosity (29%), mutations (12.2%), homozygous deletion (homodeletion; 4%), and fusion (1.2%) (Figure 1B). Among the mutations, missense mutations make up 8%, followed by 3% truncation mutations and 1.2% splice mutations (Figure 1B). Notably, we observed a significant decrease in the levels of both pre-miR-34a and mature miR-34a in *TP53* altered tumors (Figure 1C-D). Detailed dissection of *TP53* genetic alterations revealed that downregulated miR-34a expression was contributed, primarily, by heterozygous loss and missense and truncation mutations but not homodeletion (Figure 1E-F). These results, collectively, indicate that miR-34a expression is significantly reduced in PCa with TP53 abnormalities and suggest that the miR-34a based therapeutics may be particularly effective in TP53-mutated prostate tumors.

**Figure 1.**
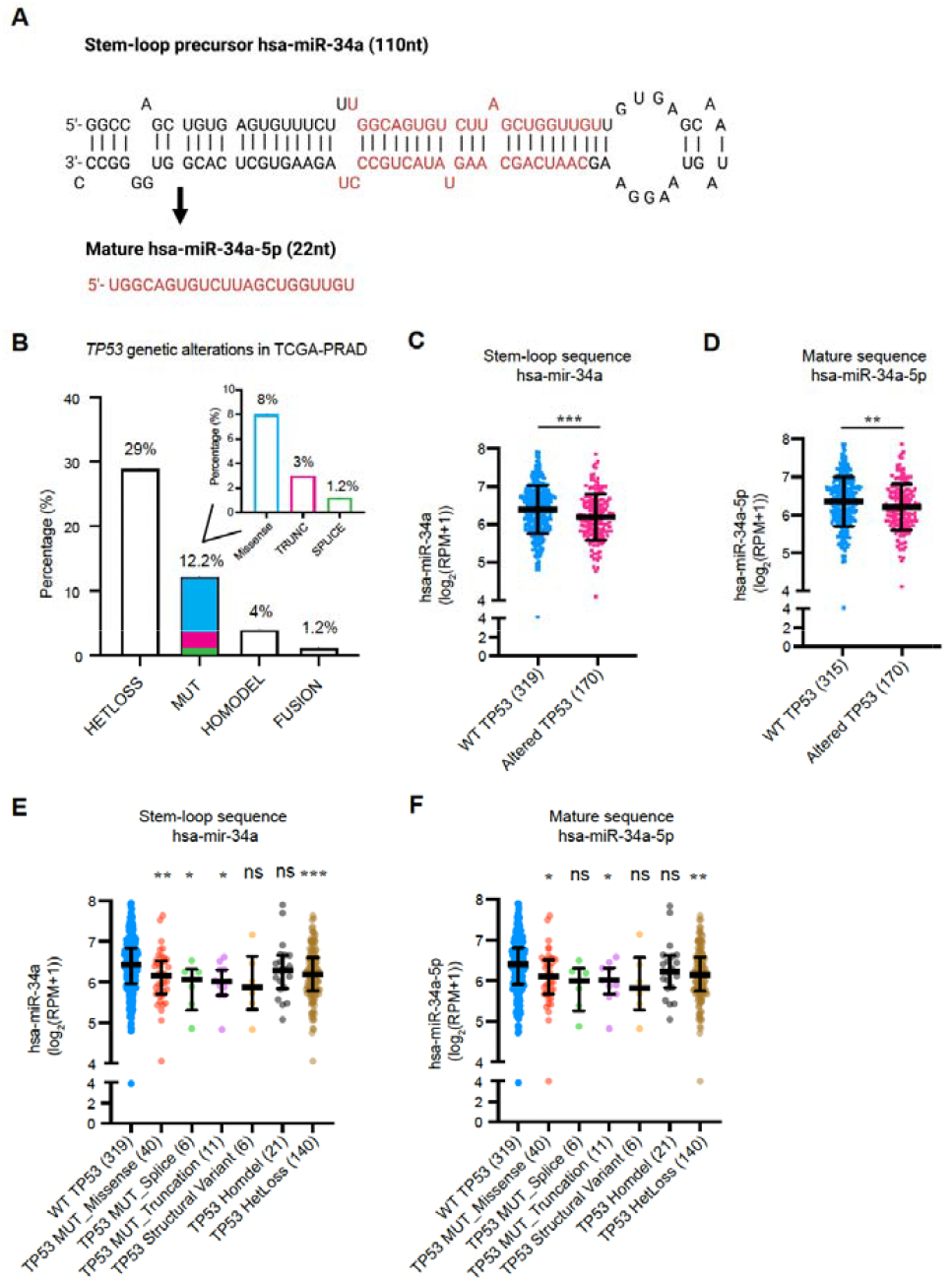
The structure of miR-34a and its regulation by TP53. **(A)** Schema of miR-34a structure (stem loop precursor and mature strand). **(B)** The frequency of *TP53* genetic alterations in PCa. The inset bar graph shows the detailed types of *TP53* mutations (MUT) including missense, truncation (TRUNC) and splice mutations. HETLoss, heterozygous loss; HOMODEL, homozygous deletion. **(C-D)** Stem loop (C) and mature (D) miR-34a expression in PCa patients with wild-type (WT) or altered *TP53*. **(E-F)** Stem loop (E) and mature (F) miR-34a expression in PCa based on *TP53* genetic alterations. All data were retrieved from cBioPortal TCGA-PRAD. ^*^, p < 0.05; ^**^, p < 0.01; ^***^, p < 0.001; ^****^, p < 0.0001; ns, not significant (Student’s t-test).

### 2.2 miR-34a mimic downregulated miR-34a targets and inhibited PCa cell growth

We employed RT-qPCR to determine the expression levels of miR-34a-5p in 4 pairs of androgen-dependent (AD) and androgen-independent (AI) human PCa xenografts (LAPC9, LNCaP, LAPC4 and VCaP), 4 cultured PCa cell lines (LNCaP, VCaP, PC3 and DU145), and one immortalized but non-transformed prostate epithelial cell line (RWPE-1). The results revealed that miR-34a was heterogeneous y expressed across xenografts and cell lines (Figure 2A). Compared to cultured RWPE-1 cells with WT *TP53*, the 4 cultured PCa cell lines showed decreased miR-34a expression (Figure 2A). Notably, miR-34a was further underexpressed in *TP53*-altered PCa cell lines (VCaP, DU145, and PC3) compared to LNCaP cells with WT *TP53* (Figure 2A). A similar trend was found in PCa xenografts where the miR-34a levels were lower in *TP53*-mutated LAPC4 and VCaP xenografts as opposed to *TP53*-WT LAPC9 and LNCaP xenografts (Figure 2A). These results are consistent with the above bioinformatics analysis showing that miR-34a expression correlates with *TP53* status (Figure 1).

**Figure 2.**
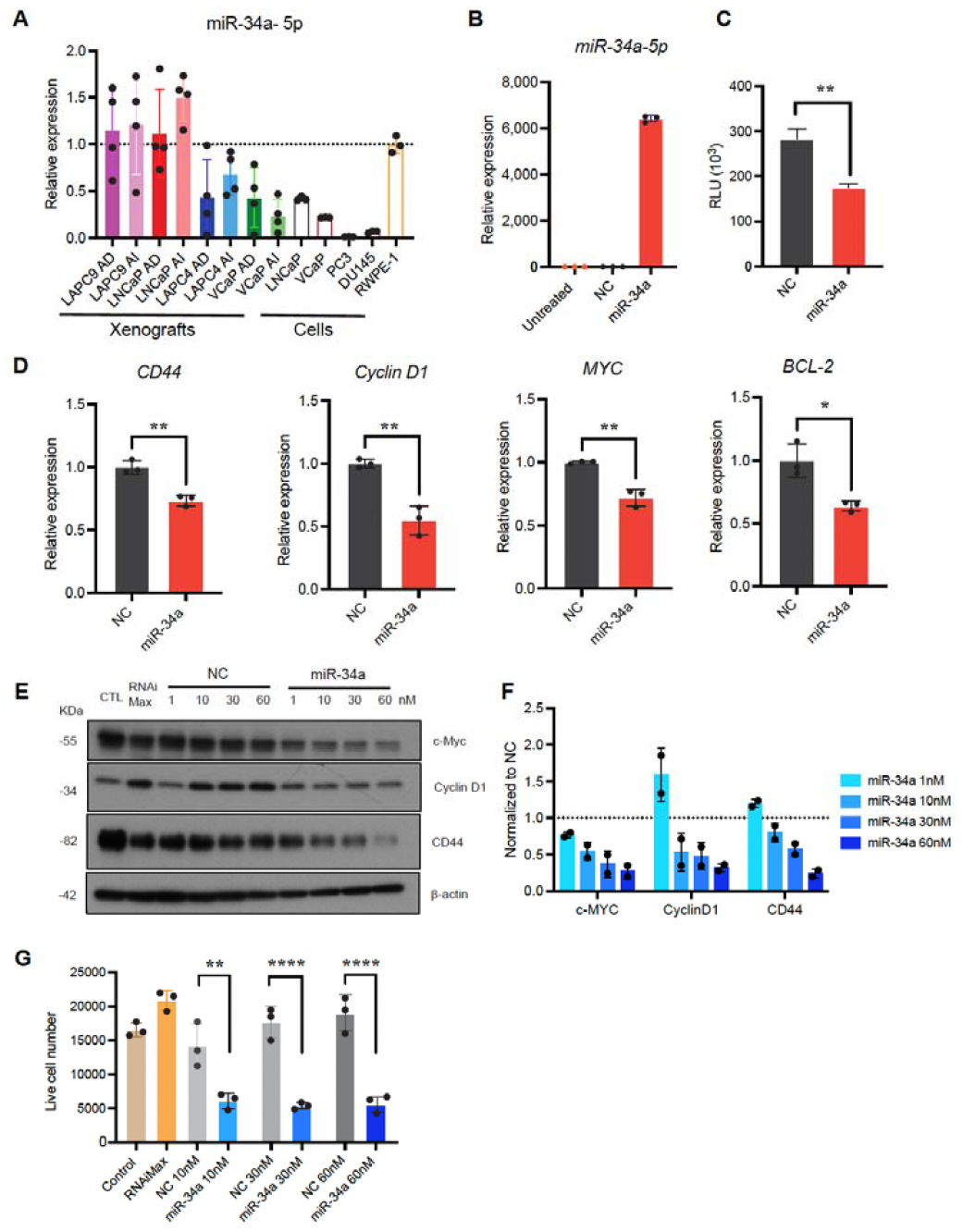
miR-34a expression in PCa cells and xenografts and miR-34a mimic downregulated molecular targets and inhibited PCa cell growth. (A) miR-34a-5p expression levels in four pairs of PCa xenografts and indicated PCa cell lines. AD, androgen dependent; AI, androgen independent. (B) miR-34a-5p expression 48 h post transfection of PC3 cells with 30 nM miR-34a mimic or NC. (n=3) (C) Targeted silencing of miR-34a Renilla sensor using miR-34a mimic in PC3-miR-34a sensor cells in vitro. Data points were normalized to NC. (n = 3) (D) Evaluation of the expression of miR-34a targets by qRT-PCR from PC3 cells following transfection with 30 nM miR-34a mimic for 48 h. (E) Representative Western blot images depicting downregulation of c-Myc, Cyclin D1, or CD44 expression following transfection of PC3 cells with miR-34a (n = 3). (F) Quantitative data from E normalized to respective NC. (G) Effect of miR-34a mimic on PC3 cell growth measured by Trypan blue cell counting 120 h after transfection of the indicated conditions. (n = 3). In all experiments, miR-34a mimic or NC were transfected into PC3 cells using lipofectamine RNAiMax. ^*^, p < 0.05; ^**^, p < 0.01; ^***^, p < 0.001; ^****^, p < 0.0001 compared to respective NC.

PC3 is the most aggressive PCa cell line being *TP53* null and AR negative and having the lowest level of miR-34a (Figure 2A). We transfected miR-34a mimic to PC3 cells using lipofectamine RNAiMax, which led to significantly increased levels of miR-34a-5p compared to PC3 cells transfected with the negative control (NC) non-targeting oligonucleotides (Figure 2B). miR-34a significantly reduced Renilla luciferase activity in PC3-miR-34a sensor cells (Figure 2C), which are PC3 cells that stably express a miR-34a Renilla luciferase (Renilla) sensor [21]. The sensor includes a miR-34a complementary sequence downstream of the Renilla luciferase gene, allowing for monitoring the targeted silencing mediated by exogenous miR-34a. Transfected miR-34a also significantly downregulated the mRNA levels of miR-34a target genes *CD44, Cyclin D1, Myc*, and *BCL-2* (Figure 2D) and protein levels of CD44, Cyclin D1 and c-Myc in a dose dependent manner (Figure 2E-F). Notably, miR-34a inhibited PC3 cell growth (Figure 2G).

### 2.3 Folate-miR-34a inhibited the growth of breast, ovarian and cervical but not PCa cells

Next, we explored the potential ‘therapeutic’ effects of folate-conjugated miR-34a, i.e., folate-miR-34a duplex, in PCa cells (Figure 3). Both miR-34a-3p passenger strand and miR-34a-5p active strand underwent partial chemical modifications with 2’ -O-methyl RNA bases (Figure 3A), and folate-miR-34a was synthesized by conjugating folate to miR-34a-3p passenger strand using click chemistry followed by an annealing step (Figure 3B). Finally, Folate-miR-34a conjugates were evaluated using polyacrylamide gel electrophoresis (Figure 3C). As the previous study has shown that folate-miR-34a is selectively targeted to breast and lung cancers overexpressing FOLR1 [21], we first confirmed the functionality of our folate-miR-34a using the MDA-MB-231 breast cancer cells as the experimental control. MDA-MB-231-miR-34a and LNCaP-miR-34a ‘sensor’ cells were established by stably expressing a miR-34a complementary sequence downstream of the Renilla luciferase gene [21]. As expected, unconjugated miR-34a duplex without transfection reagent did not show any effect in either MDA-MB-231-miR-34a or LNCaP-miR-34a sensor cells (Figure 3D-E). Seventy-two hours post transfection, folate-miR-34a significantly downregulated Renilla luciferase activity in MDA-MB-231-miR-34a sensor cells (Figure 3D) but, surprisingly, not LNCaP-miR-34a sensor cells (Figure 3E). Also, folate-miR-34a, at 200 nM, inhibited the growth of MDA-MB-231 (Figure 3F and Figure S1A), Hela (cervical cancer) (Figure 3G and Figure S1B) and OV90 (ovarian cancer) (Figure 3H and Figure S1C) cells but not that of LNCaP and PC3 PCa cells (Figure 3I-J and Figure S1D-E) although folate-miR-34a transfected by lipofectamine RNAiMax significantly repressed growth of all five cell lines at 50 nM (Figure 3F-J and Figure S1A-E). Collectively, these results indicate that folate-miR-34a, as expected, inhibited the growth of FOLR1-expressing breast, cervical and ovarian cancer cells but, unexpectedly, did not exhibit any inhibitory effects on PCa cells.

**Figure 3.**
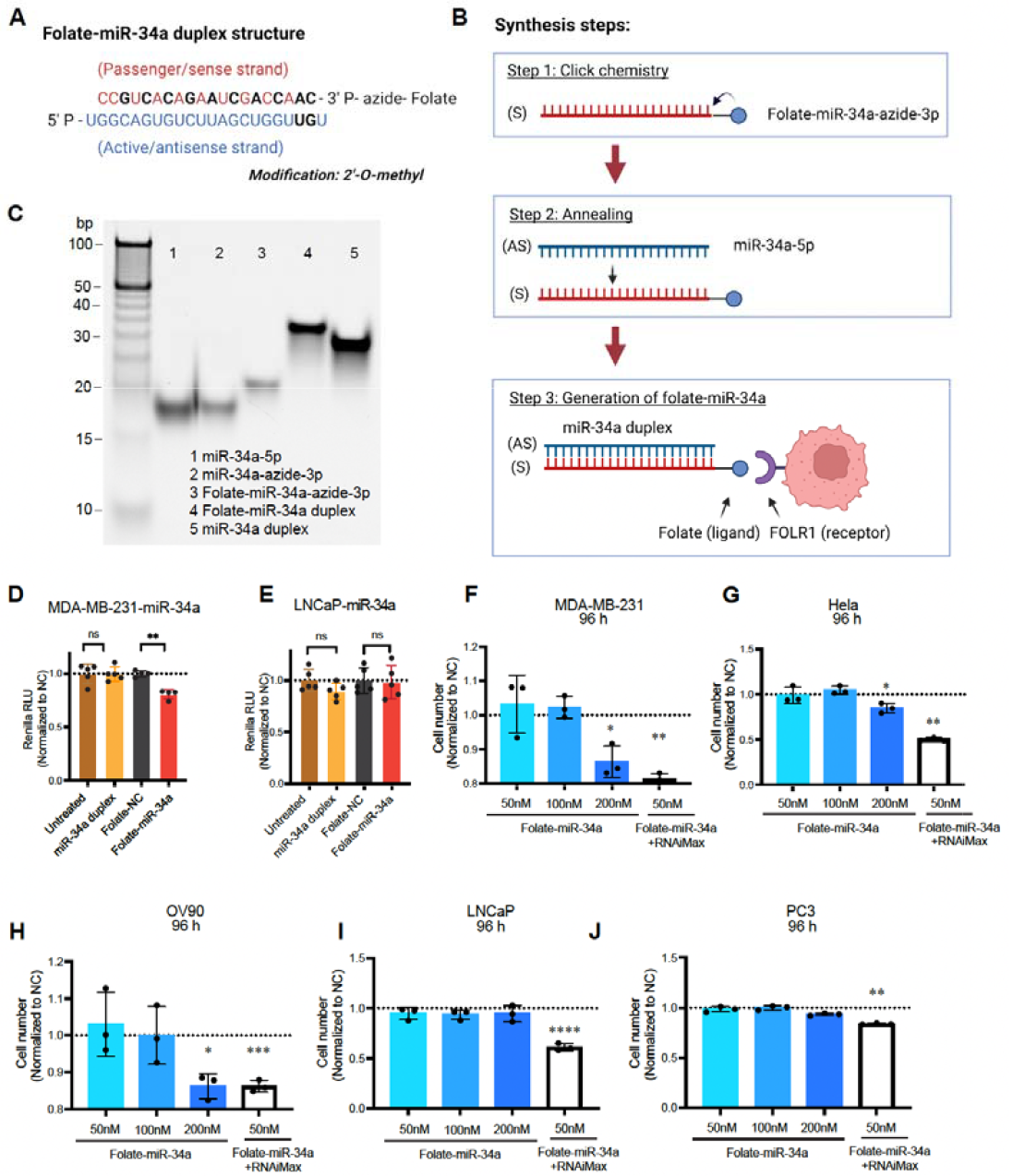
Synthesis of folate-miR-34a and its biological effects in four different cancer cells. **(A)** The structure of folate-miR-34a duplex. Chemically modified nucleotides were marked in black. **(B)** The schema of folate-miR-34a synthesis process. S, sense strand; AS, antisense strand. **(C)** Evaluation of folate-miR-34a conjugation measured by 15% native TAE PAGE. **(DE)** Targeted silencing of miR-34a Renilla sensor by folate-miR-34a in MDA-MB-231-miR-34a sensor cells (D) and LNCaP-miR-34a sensor cells (E). The results (RLU, relative luciferase unit) were normalized to folate-NC (negative control: scrambled miRNA). (n = 4) **(F-J)** Effect of folate-miR-34a on proliferation in MDA-MB-231 (breast cancer), Hela (cervical cancer), OV90 (ovarian cancer), LNCaP or PC3 cells. (n = 3). ^*^, p < 0.05; ^**^, p < 0.01; ^***^, p < 0.001; ^****^, p < 0.0001 compared to respective NC.

### 2.4 Lack of FOLR1 expression in PCa

The above results suggest that PCa cells might express little or no FOLR1. To test this possibility, we first investigated, via bioinformatics approaches, *FOLR1* mRNA levels in the normal prostate and PCa *in vivo*. The GTEx data show that among normal tissues, the levels of *FOLR1* mRNA are high in the lung, salivary gland kidney and thyroid gland but very low in the normal prostate (Figure S2A). We then evaluated *FOLR1* mRNA expression across 33 human cancers and paired normal tissues in TCGA and combined with the results in benign/normal tissues in GEPIA (Gene Expression Profiling Interactive Analysis, http://gepia.cancer-pku.cn/) database (Figure 4). We found that *FOLR1* mRNA was indeed very lowly expressed in both normal prostatic tissues and prostate tumors (PRAD) although its expression levels were high and elevated in several cancers including ovarian cancer (OV), glioblastoma (GBM), pancreatic adenocarcinoma (PAAD), rectum adenocarcinoma (READ), testicular germ cell tumor (TGCT), uterine corpus endometrial carcinoma (UCEC), and uterine carcinosarcoma (UCS) (Figure 4). On the contrary, the *FOLR1* mRNA levels were decreased in 6 cancers including breast cancer (BRCA), lung squamous cell carcinoma (LUSC), head and neck squamous cell carcinoma (HNSC), kidney chromophobe (KICH), acute myeloid leukemia (LAML), and skin cutaneous melanoma (SKCM) (Figure 4). Interestingly, the *FOLR1* mRNA levels, although low, were also reduced in PCa compared to normal prostate (Figure S2B). Intriguingly, the low levels of *FOLR1* mRNA were detected preferentially in AR^+^ luminal epithelial cells in normal human prostate (Figure S2C) based on our RNA-seq analysis using purified cell populations [30]. Also, the low *FOLR1* mRNA levels showed increased tendency on two PCa patient cohorts [31,32] who went through short-term neoadjuvant androgen-deprivation therapy (ADT) (Figure S2D-E).

**Figure 4.**
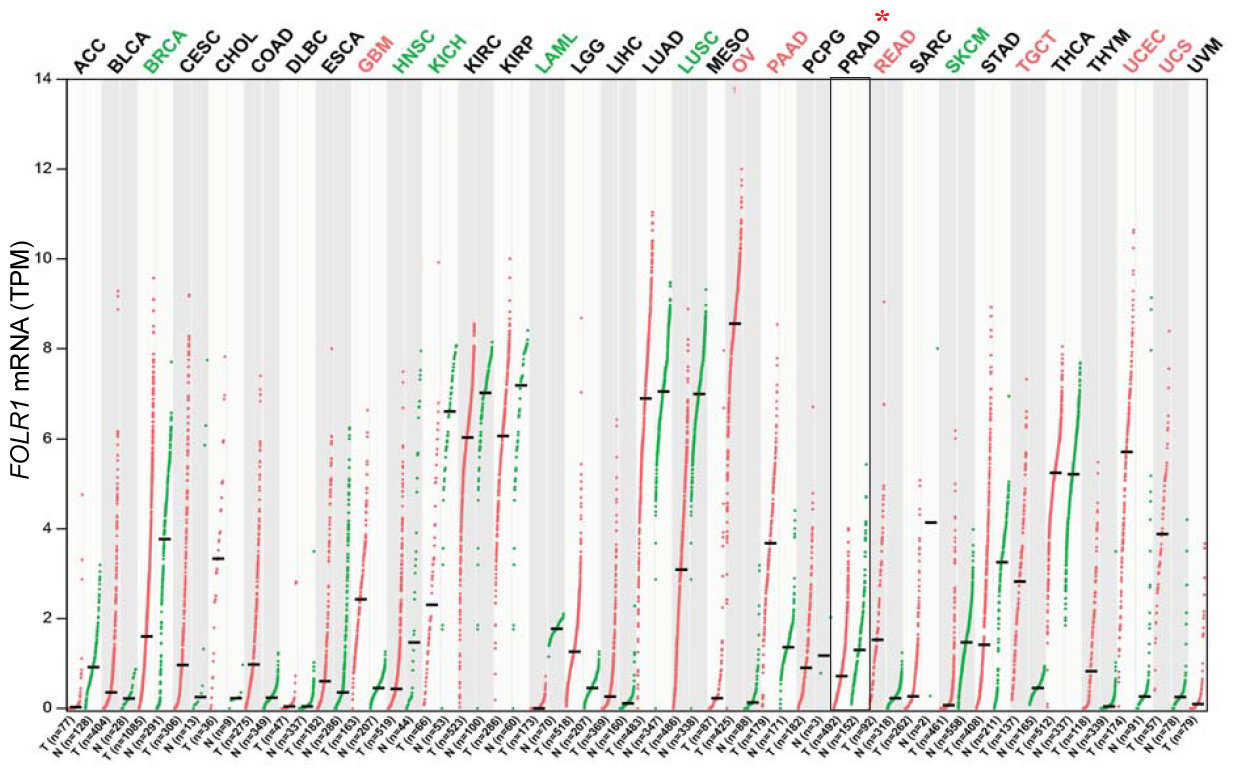
FOLR1 mRNA expression across 33 human tumors (T) and paired normal tissues (N). Significant increase or decrease in T compared to N is highlighted in red or green respectively.

Next, we investigated expression of FOLR1 mRNA and protein levels in PCa cell lines and xenografts (Figure 5). In CCLE (Cancer Cell Line Encyclopedia), *FOLR1* mRNA was barely expressed in PCa cell lines in contrast to ovarian, endometrial and kidney cancer cells that highly expressed *FOLR1* mRNA (Figure S2F and Figure 5A). Our RT-qPCR (reverse transcription – quantitative polymerase chain reaction) and Western blotting also revealed FOLR1 to be barely expressed in PCa cells as well as prostate xenograft tumors at both mRNA and protein levels (Figure 5B-C). We also assessed the FOLR1 expression in PC3, LNCaP and Hela (positive control) cells using immunofluorescence (Figure 5D) and flow cytometry (Figure 5E-G), both of which showed that PC3 and LNCaP cells did not express FOLR1. To further visualize folate uptake *in vitro* and *in vivo*, we conjugated a near-infrared (NIR) dye to folate (i.e., folate-NIR). As shown in Figure 5H-J, in contrast to Hela cells, LNCaP and PC3 PCa cells did not show any folate-NIR uptake in the cells.

**Figure 5.**
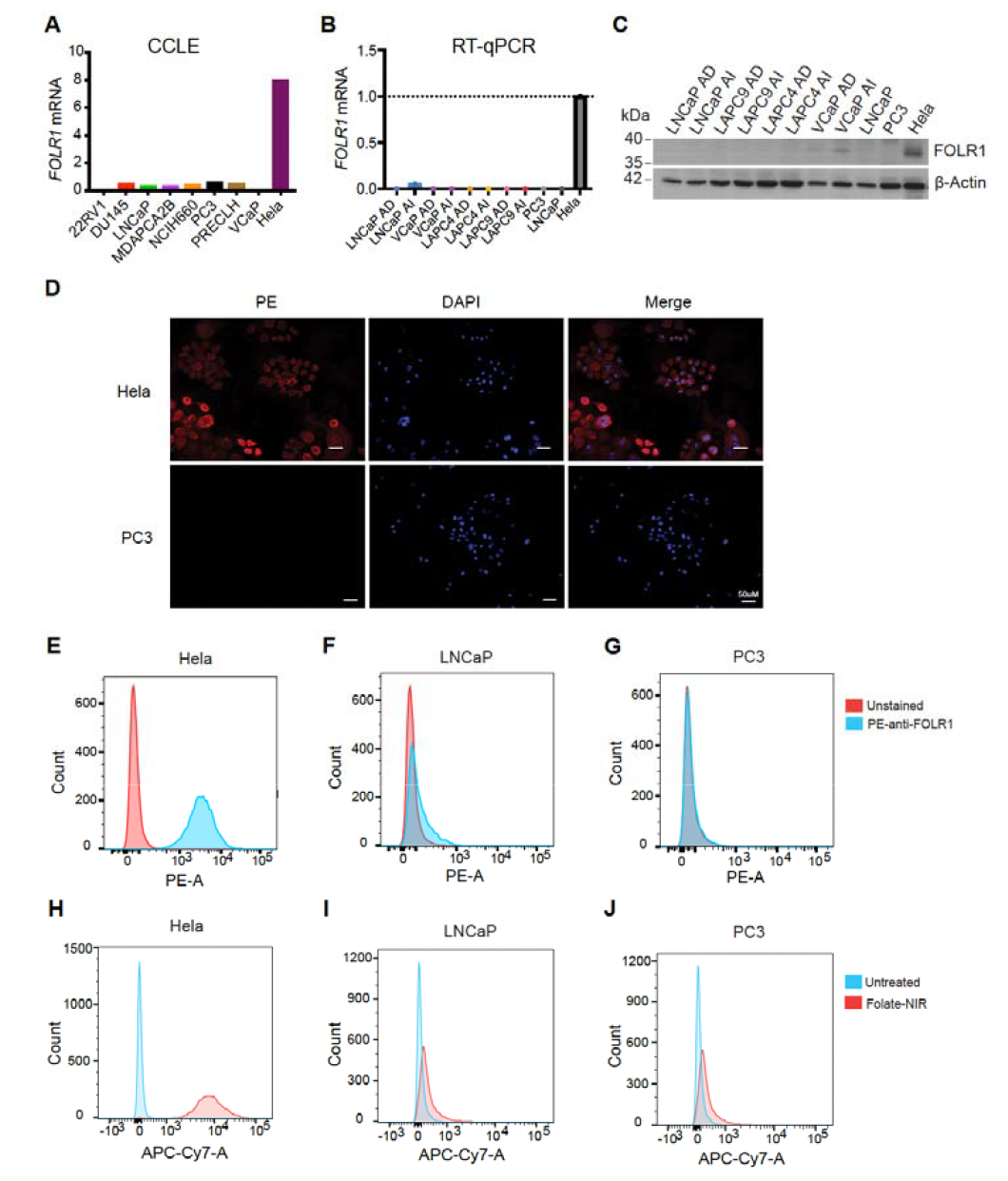
FOLR1 expression in PCa. (A) FOLR1 expression in PCa cell lines in the Cancer Cell Line Encyclopaedia (CCLE). (B-C) The mRNA (B) and protein (C) levels of FOLR1 in four pairs of PCa xenografts and cell lines. (D) Representative immunofluorescent images showing the expression of FOLR1 (PE) in Hela cells but not in PC3 cells. (E-G) Flow cytometry analysis (using PE-conjugated anti-FLOR1 antibody) showing expression of FOLR1 in Hela (E) but not in LNCaP (F) or PC3 (G) cells. (H-J) Folate-NIR uptake in FOLR1+ Hela cells (H) compared to FOLR1-LNCaP (I) and PC3 (J) cells. Histograms represent overlaid flow cytometry data of unstained cells and cells stained with folate-NIR (50 nM) Flow cytometry was analyzed on Cy7 channel.

### 2.5 Folate-miR-34a also did not accumulate nor show any effect in PSMA-expressing PC_a_ cells

The above results indicate that folate-miR-34a may not be an effective PCa-targeting therapeutic due to lack of appreciable FOLR1 expression. Folate (folic acid) has been reported to bind to another molecule, prostate specific membrane antigen (PSMA) [33,34]. PSMA, also known as glutamate carboxypeptidase II, is a type II membrane protein that is highly expressed in PCa [35]. PSMA is constitutively internalized and rapidly recycles back to cell surface enabling additional rounds of internalization [19,36,37]. Yao et al. reported that PSMA can bind to folate at pH 7.4 and functions as a folate transporter and that PSMA expression significantly increased cellular uptake of folic acid under conditions of limiting folate in PCa cells [34]. Moreover, folate was able to compete with other substrate and inhibit the enzymatic activity of PSMA [34], indicating the folate-binding capability of PSMA. These studies suggest that folate-miR-34a might be able to gain access to PSMA-expressing PCa cells.

Unfortunately, we did not observe any biological effects of folate-miR-34a on PSMA-expressing LNCaP cells (Figure 3E and I; Figure S1D) nor did we observe folate-NIR uptake in LNCaP cells (Figure 5I). In addition, fluorescence microscopy studies revealed prominent Folate-NIR accumulation in FOLR1-expressing Hela cells but not in PC3 cells, which do not express FOLR1 or PSMA nor in LNCaP cells, which lack FOLR1 but do express PSMA [38] (Figure 6A-C). We also conducted in vivo biodistribution studies of Folate-NIR to monitor its targeting specisficity in PCa xenografts, and the results revealed that 24 h after injection, Folate-NIR was not significantly retained in LNCaP AD/AI or LAPC9 AD/AI tumor tissues but cleared mainly by the kidney of the host mice (Figure 6D-G), whereas it was significantly accumulated in Hela tumor tissues (Figure 6H). Analysis of tumor/kidney ratio further supported limited accumulation of folate-NIR in prostate tumors as compared to Hela tumor that highly expresses FOLR1 (Figure 6I).

**Figure 6.**
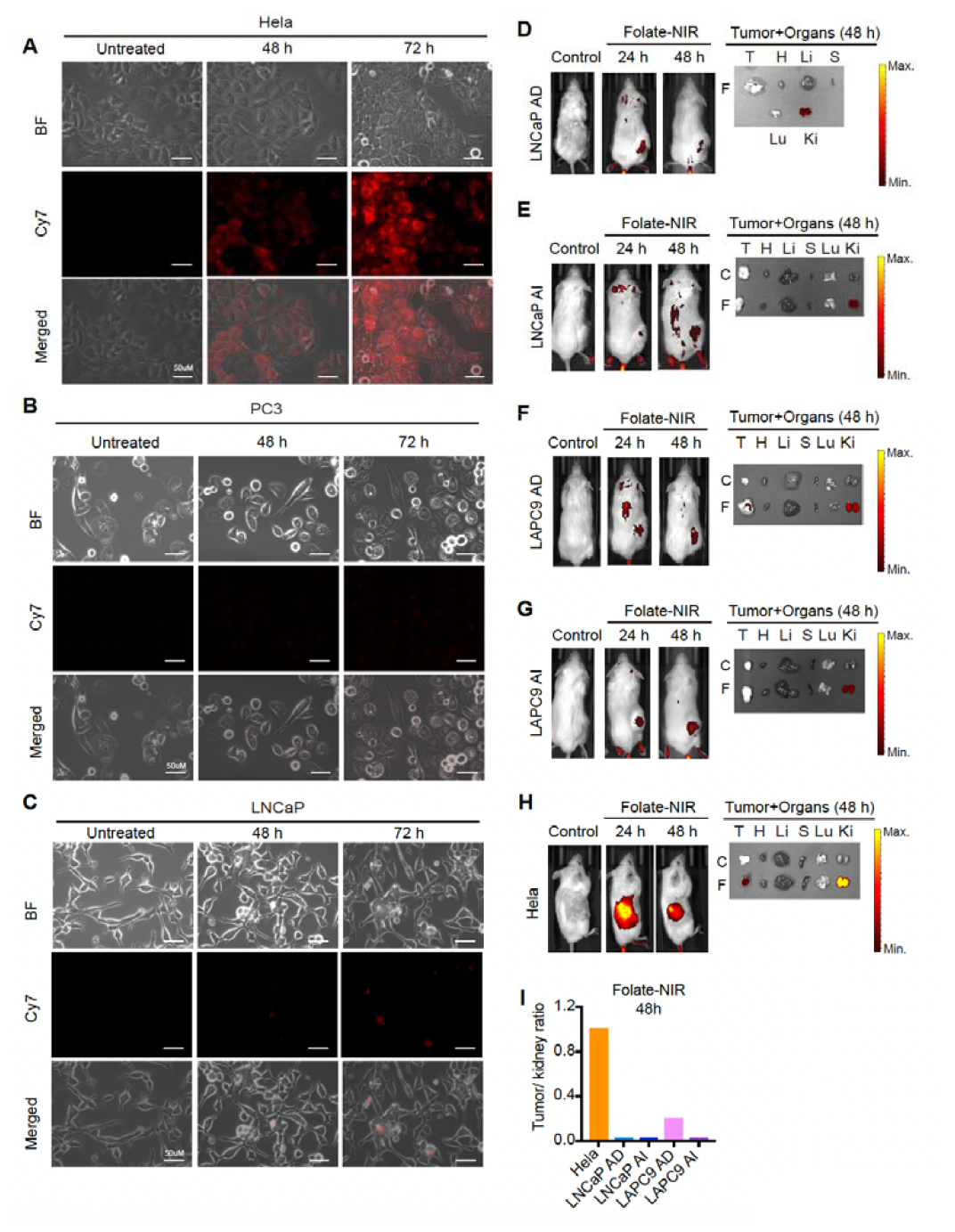
*In vitro* cellular uptake and *in vivo* biodistribution of folate-NIR in PCa. (A-C) Immunofluorescence analysis of Folate-NIR uptake in FOLR1+PSMAHela cells (A) compared to FOLR1-PSMA-PC3 (B) and FOLR1-PSMA+ LNCaP (C) cells. Scale bars, 50 μm. Folate-NIR was analyzed using Cy7 filter. (D-H) Left: Representative live imaging of mice bearing the indicated LNCaP AD/AI, LAPC9 AD/AI, or Hela xenograft tumors after intravenous injection of a single dose (10 nmol) of folate-NIR and analyzed at indicated time points (Control, time 0). Right: Ex vivo tissue biodistribution in mice 48 h after administering folate-NIR (T, tumor; H, heart; Li, liver; S, spleen; Lu, lungs; Ki, kidneys). (I) The tumor/ kidney ratio determining folate-NIR retention in xenograft tumors quantified from D-H.

Collectively, these data suggest that folate is not a suitable ligand for targeting miR-34a efficiently to PCa cells.

## 3. Discussion

Many studies have demonstrated that miR-34a represents a promising anti-CSC inhibitor for treating advanced PCa [5,7,11]. However, lack of efficient and safe delivery strategies remains a major bottleneck in the clinical failed in large part due to systemic toxicity caused by the packaging vehicle and immunotoxicity likely associated with miR-34a over-dosing [7,39,40]. In this study, we attempted to develop a package-free targeted delivery platform, i.e., folate-miR-34a, for PCa therapy. Unfortunately, folate-miR-34a did not exhibit appreciable uptake in PCa cells and did not elicit any PCa-inhibitory effects due to lack of expression of FOLR1, the major high-affinity receptor of folate, in PCa cells. Folate-miR-34a did not even show any uptake and biological effects in PCa cells that express PSMA, which can bind folate (as an enzymatic substrate) and function as a folate transporter [34,41]. In PC3 cells that do not express endogenous PSMA, PSMA re-expression significantly increased cellular uptake of folic acid under conditions of limiting folate [34]. Also, folate was able to compete with other substrates and inhibited the enzymatic activity of PSMA [34], further supporting the folate-binding capability of PSMA. Furthermore, previous studies reported that folate-conjugated delivery platforms could achieve specific delivery of various payloads to PSMA-expressing PCa cells and demonstrated better anti-cancer efficacy *in vivo* as compared to non-targeted ones [38,42-45]. These early studies [34,38,41-45] provide the rationale for targeted delivery of folate-conjugated miR-34a to PSMA-expressing PCa. However, our data herein show that folate-miR-34a was not uptaken by and did not achieve specific delivery to PSMA^+^ LNCaP cells. These ‘inconsistent’ results may likely be related to different chemistries of folate-drug conjugates, and a significant difference is that all previous studies conjugated folate to liposomes, nanocarriers or bacterial minicells. Since the payloads are encapsulated in those vehicles, drugs may get internalized into the PCa cells through endocytosis, and the observed therapeutic effects, in principle, could be due to off-target (i.e., PSMA-independent) effects. Another consideration is that the binding affinities for folate to FOLR1 and PSMA are different. The affinity of folate for PSMA is much lower than FOLR1 [19], which could result in less miR-34a being delivered. Also, very little miR-34a was released from endosome due to endosome entrapping. These two limitations could together lead to minimal therapeutic effect of folate-miR-34a on PSMA-expressing PCa cells. On the other hand, folate-miR-34a inhibited cell proliferation in breast, cervical and ovarian cancer cells that highly express FOLR1, indicating its potential therapeutic applications for these FOLR1 expressing cancers. FORL1 has been associated with tumor relapse and chemotherapy resistance in cervical and ovarian cancers [46,47]. FOLR1 has emerged as an optimal target and multiple FOLR1 targeting therapeutic approaches have been or are being tested preclinically and clinically [48-52]. In-depth preclinical studies of folate-miR-34a are needed to validate the efficacy and safety in cervical and ovarian cancers.

PSMA is highly expressed in metastatic CRPC and has been shown to be a validated therapeutic target. Currently ARX517, an antibody-drug conjugate composed of a fully humanized anti-PSMA mAb linked to AS269 as a potent microtubule inhibitor, is being studied in a phase I/II clinical trial enrolling patients with metastatic CRPC. DUPA is a synthetic urea-based ligand that can bind to PSMA with high affinity leading to saturation of the receptor in a short period of time [53]. Thomas et al. were the first to use DUPA conjugates to deliver siRNAs selectively to PSMA-expressing PCa cells [54]. Treating LNCaP cells with fluorescently tagged siRNA directly linked to DUPA (DUPA-siRNA-cy5) in vitro resulted in substantial uptake within 1 h of treatment [48]. Similarly, significant accumulation of DUPA-siRNA was observed in LNCaP xenograft tumors after intravenous injection of DUPA-siRNA-cy5. Another study by Tai et al. showed that DUPA-siRNA induced tumor growth inhibition in LNCaP xenografts [55]. These studies suggests that DUPA-conjugated miR-34a could be a potential therapeutic to target PSMA-expressing PCa. Considering the huge success of lipid nanoparticle (LNP) application in COVID-19 vaccines, DUPA-conjugated LNP with miR-34a as payload could be another novel delivery approach to explore in the future.

Ligand-directed miR-34a delivery potentially represents a novel strategy to achieve specific and efficient delivery for targeting a wide range of cancer types. In comparison to packaged vehicles, this strategy circumvents off-target effects and non-specific biodistribution that result in systematic toxicity. Nevertheless, there are still many obstacles to overcome in translating the ligand-directed miR-34a as PCa-targeting therapeutics (Figure 7). For example, PCa lack appreciable FOLR1 expression invalidating folate-miR-34a as a therapeutic in this cancer (Figure 7A). Another concern is endosomal entrapment of ligand-conjugated miR-34a (Figure 7B), which, in fact, represents the major challenge for ligand-conjugated miRNA delivery. Once ligand-conjugated miR-34a is internalized in PCa cells, miR-34a must successfully escape from the endosome into the cytoplasm, where they can interact with the RNAi machinery. Otherwise, it will be subject to lysosomal degradation when the late endosomes fuse with the lysosomes. Various strategies have been developed to promote endosomal release including cell-penetrating peptides, fusogenic and endolytic peptides, or chemical agents such as chloroquine and nigericin [56-60]. Some of these approaches are limited in translation due to systematic toxicity *in vivo*, and more efforts are needed to develop effective and less toxic endosomal agents aiming to further enhance the efficacy of ligand-conjugated miR-34a therapeutics.

**Figure 7.**
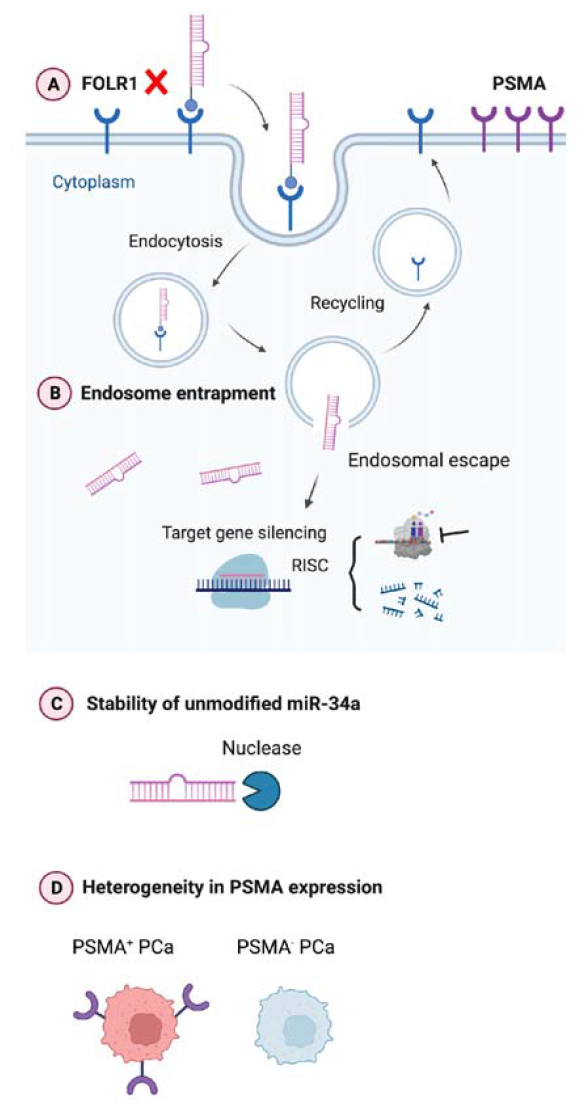
Graphical abstract. Current challenges in developing ligand-directed miR-34a therapeutics in PCa. See Text for details.

Instability and (exo)nuclease-mediated degradation of unmodified miR-34a presents another issue (Figure 7C). Chemical modifications of miRNAs, including introduction of 2′-O-methyl and 2′-fluoro to the ribose, and phosphorothioate substitutions to the backbone, have been used to improve serum stability and increase intracellular half-life to ultimately reduce the therapeutic doses [19,61]. miR-34a used in current study is a partially modified (PM) version containing a minimal number of 2′-O-methyl modifications akin to commercially available miR-34a mimics [21,62]. In preliminary studies, we observed that both PM-miR-34a duplex and PM-folate-miR-34a duplex started degradation from 10 min and nearly 50% oligos were degraded within 30 min, demonstrating the instability of PM-miR-34a oligos (Figure S3). This suggests that chemical modifications should be carefully designed and selected to improve the stability of miR-34a therapeutics. A very recent study by Abdelaal et al. demonstrated that full chemical modification of miR-34a duplex enhances both stability and activity of miR-34a [62]. This improved stability could be beneficial for *in vivo* applications by reducing the effective dose as well as minimizing toxicity resulting from higher miRNA doses or administration frequency.

Finally, the heterogeneity in expression levels of targets, i.e., cancer cell surface receptors (e.g., FOLR1) and membrane proteins (e.g., PSMA) should be considered as one of the limiting factors for ligand-directed miR-34a therapeutics (Figure 7D*). A priori*, PSMA represents an ideal target for PCa treatment due to its highly specific cell surface expression, which lends possibilities for both imaging and therapeutic development. However, PSMA expression exhibits significant intra- and inter-tumor heterogeneity in advanced PCa [63-65], as also highlighted by recent preclinical and clinical studies demonstrating that not all patients with PSMA-positive PCa respond to PSMA-targeted radionuclide ^177^Lu-PSMA-617 [66]. PSMA expression is known to be (partially) regulated by AR, and AR^+^ PCa cells are generally PSMA^+^ whereas AR^-^ PCa cell lines are PSMA^-^. But interestingly, a subset of AR^-^ CRPC and neuroendocrine PCa (NEPC) did express PSMA [63], indicating alternative mechanisms of PSMA regulation including, among others, transcriptional regulation by HOXB13 [63], deleterious DNA repair aberrations [64], and epigenetic silencing by CpG methylation [65]. Some pharmacological approaches can augment PSMA levels in PCa. For example, treatment with histone deacetylase (HDAC) inhibitors reversed the epigenetic suppression, leading to PSMA re-expression both *in vitro* and *in vivo* [65]. A deeper understanding of the biology of PSMA in PCa is needed to elucidate pharmacological strategies that can help tackle the heterogeneity in PSMA expression (Figure 7D) and thus enhance the efficacy of PSMA-targeted therapies for advanced PCa.

## 4. Materials and Methods

### 4.1. Cell lines and animals

LNCaP, PC3, DU145, VCaP, and RWPE-1 cells were purchased from the American Type Culture Collection (ATCC). MDA-MB-231-miR-34a reporter cells and LNCaP-miR-34a sensor cells were kind gifts from Dr. Andrea Kasinski (Purdue University). PC3-miR-34a sensor cells were generated as described previously [21]. MDA-MB-231, OV90 and Hela cells were gifts from Drs. Chetan Oturkar and Shamshad Alam (Roswell Park Comprehensive Cancer Center). VCaP cells were cultured in DMEM medium supplemented with 10% fetal bovine serum (FBS) and antibiotics. RWPE-1 were cultured in Keratinocyte Serum Free Medium (K-SFM) supplemented with 0.05 mg/ml bovine pituitary extract (BPE) and 5 ng/ml human recombinant epidermal growth factor (EGF). Except VCaP and RWPE-1 cells, all other cell lines were cultured in RPMI medium plus 10% heat-inactivated fetal bovine serum (FBS) plus antibiotics. These cell lines were authenticated regularly in our institutional CCSG Cell Line Characterization Core and also examined to be free of mycoplasma contamination. LAPC4 and LAPC9 xenograft lines were initially provided by Dr. Robert Reiter (UCLA) and have been used extensively in our previous studies [4,11]. Immunodeficient mice, NOD/SCID (non-obese diabetic/severe combined immunodeficiency) and NOD/SCID-IL2Rγ™/™ (NSG) were obtained from the Jackson Laboratory, and breeding colonies were maintained in standard conditions in our animal facilities.

### 4.2 Preparation of folate-miR-34a duplex

miR-34a duplex was constructed using two RNA oligonucleotides: miR-34a-5p guide strand and miR-34a-3p passenger strand (Integrated DNA Technologies). A scrambled miRNA (negative control, NC) synthesized with the same modifications was used to form a control duplex. The synthesis of folate-miR-34a duplex was previously described [21]. In brief, a click reaction was conducted between folate-DBCO and azide-modified passenger miR-34a (or scramble). Click reaction was performed at a 1:10 molar ratio (azide oligo/folate-DBCO) at room temperature in water for 10 hours and then cooled to 4°C for overnight. Unconjugated folate was removed from the reaction using Oligo Clean & Concentrator (Zymo Research) per the manufacturer’s instructions.

After conjugation, the miR-34a-5p guide strand was annealed to the folate-miR-34a-azide-3p passenger strand at an equal molar ratio in the presence of annealing buffer [10 mM Tris buffer, pH 7 (Sigma), 1 mM EDTA (Sigma), 50 mM NaCl (Sigma)] followed by incubation at 95 °C for 5 min, and slow cooling to room temperature for 1.5 hours. Annealed oligos were then stored at ™80 °C. Conjugation was verified using 15% tris base, acetic acid, EDTA (TAE) native PAGE.

### 4.3 RNA isolation and Real-time RT-PCR analysis

Total RNA was extracted using Direct-zol RNA Miniprep Plus Kits (Zymo Research) according to the manufacturer’s instructions. RNA concentration was quantified using a nanodrop. qRT-PCR was performed using a CFX Connect Real-Time PCR Detection System (Bio-Rad). For miR-34a-5p expression assay, qPCR data were normalized to U6. For miR-34a target genes, qPCR data were normalized to GAPDH. Data were then analyzed using the 2™ΔΔCt method and expressed as fold change.

### 4.4 In vitro Renilla Luciferase assay

MDA-MB-231-miR-34a reporter cells or LNCaP-miR-34a sensor cells were treated with 100nM miR-34a duplex, 100nM folate-NC, and 100nM folate-miR-34a (hereafter referred to as folate-34a). At 72h, Renilla-Glo Luciferase assay (Promega) was performed as per manufacture instructions. In brief, Renilla-Glo Luciferase substrate was mixed with Renilla-Glo buffer at 1:1000 dilution followed by addition into each well. After shaking the plates at room temperature for 10 min, Renilla luciferase signal was measured using a BioTek Synergy microplate reader (Agilent).

### 4.5 Cell proliferation assays

For the functionality validation of miR-34a mimic (MC11030, Thermo Fisher), PC3 cells were seeded onto individual wells of a 96-well plate. The next day, cells were transfected with miR-34a or NC at the indicated concentrations using Lipofectamine RNAiMAX (Life Technologies). At 96h, Trypan blue excluding cells were counted with a hemocytometer then compared to coresponding NC to determine relative cell growth. Similarly, for the evalution of effects of folate-miR-34a in multiple cancer cell lines, cells were seeded onto individual wells of a 96-well plate. The next day, cells were treated with folate-miR-34a at the indicated concentrations, or transfected with 50nM folate-miR-34a using Lipofectamine RNAiMAX (Life Technologies). At the indicated time points, Trypan blue excluding cells were counted with a hemocytometer to determine relative cell growth by comparing to coresponding NC.

### 4.6 Western blotting

For Western blotting analysis, whole cell lysate was prepared in RIPA buffer and run on 4–15% gradient SDS-PAGE gels. The proteins were transferred to nitrocellulose membrane followed by incubation with primary antibodies and corresponding secondary antibodies. Films were developed using Western Lighting ECL Plus reagent (PerkinElmer). Antibodies used: c-Myc (E5Q6W) rabbit mAb (18583, Cell Signaling), Cyclin D1 (E3P5S) XP rabbit mAb (55506, Cell Signaling), CD44 (156-3C11) mouse mAb (3570, Cell Signaling), and FOLR1 mouse mAb (sc-515521, Santa Cruz).

### 4.7 Immunofluorescence (IF)

Hela and PC3 cells were seeded onto 6-well plate, each containing a sterilized coverslip. On the next day, medium was aspirated and 1 ml 4% formaldehyde was added to each well for 10 min. Coverslips were rinsed with 1X DPBS 3 times for 5 min each. Then cells were blocked with 5% BSA solutions at room temperature for 1h. Then blocking solution was aspirated and the cells were incubated with phycoerythrin (PE) anti-FOLR1 antibody (908304, Biolegend) at room temperature for 1h. Coverslips were rinsed with 1X DPBS followed by mounting with ProLong Gold Antifade Reagent with DAPI (P36931, Thermo Fisher).

### 4.8 Flow cytometry

FOLR1^+^ Hela cells, FOLR1^™^ LNCaP cells, and FOLR1^™^ PC3 cells growned in 10 cm culture dish were harvested (cell viability > 90%) and washed three times in ice-cold DPBS buffer supplemented with 0.5% BSA and aliquoted to a density of 5× 10^6^ cells/ml in Flow Cytometry Staining Buffer (FC001, R&D Systems). Next, flow cytometric analyses were performed following standard protocols. In brief, 5 × 10^5^/100ul cells were incubated with Fc Receptor Binding Inhibitor Polyclonal Antibody (14-9161-73, eBioscience) on ice for 20 minutes. Without washing, staining was proceeded with primary antibody PE anti-FOLR1 antibody (908303, BioLegend) incubation followed by flow cytometric analysis using LSRFortessa flow cytometer (BD Biosciences). Data were analyzed using FlowJo software v10 (Tree Star Inc.). For the functionality of FOLR1, Hela cells, LNCaP cells, and PC3 cells were incubated with Folate-NIR (50 nM) for the indicated time peroids followed by flow cytometric analyses as described above.

### 4.9 Fluorescence microscopy

Hela cells, LNCaP cells, and PC3 cells were seeded into 6-well culture plate. Then spent medium were replaced with fresh medium containing Folate-NIR (50 nM). At the indicated time points, fluorescence images were acquired using an KEYENCE BZ-X All-in-One Fluorescence Microscope.

### 4.10 Whole body imaging and tissue biodistribution

Xenograft model: Briefly, all PCa AD (androgen-dependent) and AI (androgen independent) xenograft tumors (LNCaP and LAPC9) were routinely maintained in intact immunodeficient NOD/SCID or NSG mice [4,11]. For the AI lines, parental AD tumor cells were purified, mixed with Matrigel, injected subcutaneously and serially passaged in surgically castrated immunodeficient mice. For Hela xenograft tumors, Hela cells (2 × 10^6^) were injected into the flank of 6–8 week-old female NOD/SCID mice. Once tumors reached approximately 300-400 mm^3^ in volume, animals (2 mice/ group) were intravenously injected with 10 nmol of Folate-NIR in PBS. For vehicle control group, PBS was intravenously injected with PBS. Whole-body, fluorescence images were acquired in vivo at indicated time points using IVIS^®^ Spectrum (Ex/Em = 745/800 nm). After whole-body imaging at the end point, animals were dissected and selected tissues were analyzed for fluorescence intensities using the IVIS^®^ imager. The tumor-to-kidney ratio was calculated by dividing the average radiant efficiency of tumor by the average radiant efficiency of kidney for each xenograft.

### 4.11 Statistical analysis

Statistical analysis was performed using Prism statistical software. Unpaired two-tailed Student’s t-test was used to compare significance between two groups. One-way ANOVA was used to compare the differences among multiple groups and multiple comparisons were corrected using Tukey’s post hoc test. The results were presented as mean ± S.D as denoted in the figure legends. Statistically significant p-values are as indicated in the corresponding figure legends.

## 5. Conclusions

- Folate-miR-34a did not elicit PCa-inhibitory effects due to a lack of appreciable expression of FOLR1 in PCa cells.
- Folate-miR -34a also did not display any apparent effect on PCa cells expressing prostate-specific membrane antigen (PMSA) despite the reported folate’s binding capability to PSMA.
- Folate-miR-34a exhibited impressive inhibitory effects on breast, ovarian and cervical cancer cells, suggesting its potential therapeutic application on FOLR1-expressing cancers including ovarian and cervical cancers.
- The insights offered from the current study highlight challenges in specific delivery of folate-miR-34a to PCa due to lack of target receptor expression and shed lights on the future development of ligand-conjugated miR-34a as potential therapeutics for advanced and aggressive PCa.

## Supplementary Materials

The following supporting information can be downloaded at: www.mdpi.com/xxx/s1.

## Author Contributions

W.L. conceived the project with D.G.T., designed and performed most experiments, interpreted the results, and wrote the manuscript draft. Y.W. provided partial bioinformatic data on FOLR1 and performed IF and some of qPCR experiments. X.L. generated bioinformatics data on TP53. S.W. performed partial cell proliferation experiment. M.W. performed western blotting experiment. S.G.T. and J.A.S. offered technical support on the IVIS imaging. A.T. maintained and provided human prostate cancer xenografts. A.M.A. and S.K. provided folate-conjugates and key reagents. I.P., G.C. and A.L.K. discussed the content and critically reviewed the manuscript draft. D.G.T. aided in manuscript writing and finalized the manuscript. All authors have read and agreed to the published version of the manuscript.

## Institutional Review Board Statement

All animal-related studies in this study have been approved by our Institutional Animal Care and Use Committee (IACUC) at the Roswell Park Comprehensive Cancer Center (animal protocol# 1331 M and 1328 M).

## Informed Consent Statement

Not applicable.

## Data Availability Statement

The datasets used and/or analyzed in the study are available from the corresponding authors upon request.

## Acknowledgments

We would like to thank the Translational Imaging Shared Resource in Roswell Park Comprehensive Cancer Center for the support (IVIS S10 grant S10 OD16450 and CCSG core grant P30CA016056).

---

**Figure S1.**
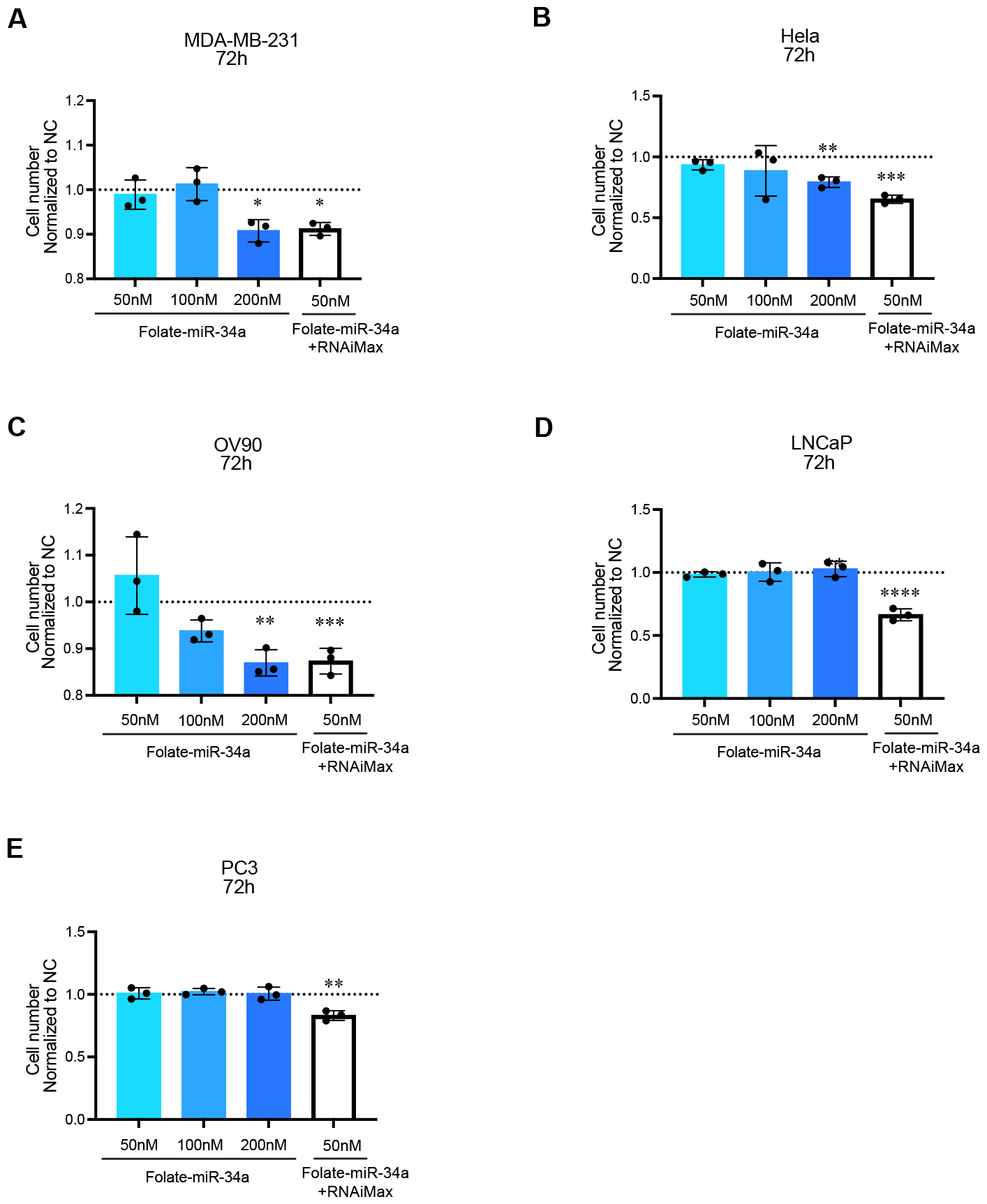

**Figure S2.**
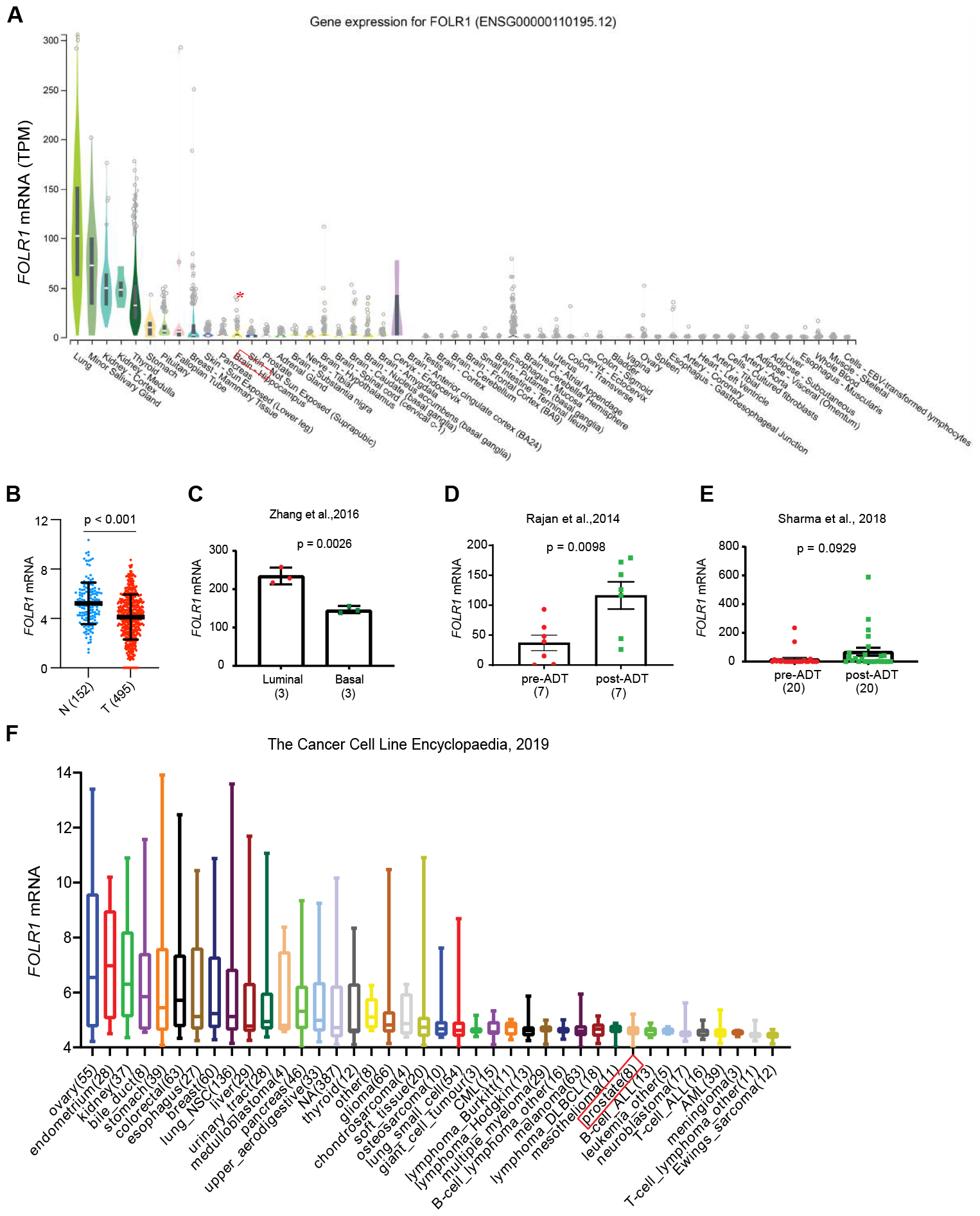

**Figure S3.**
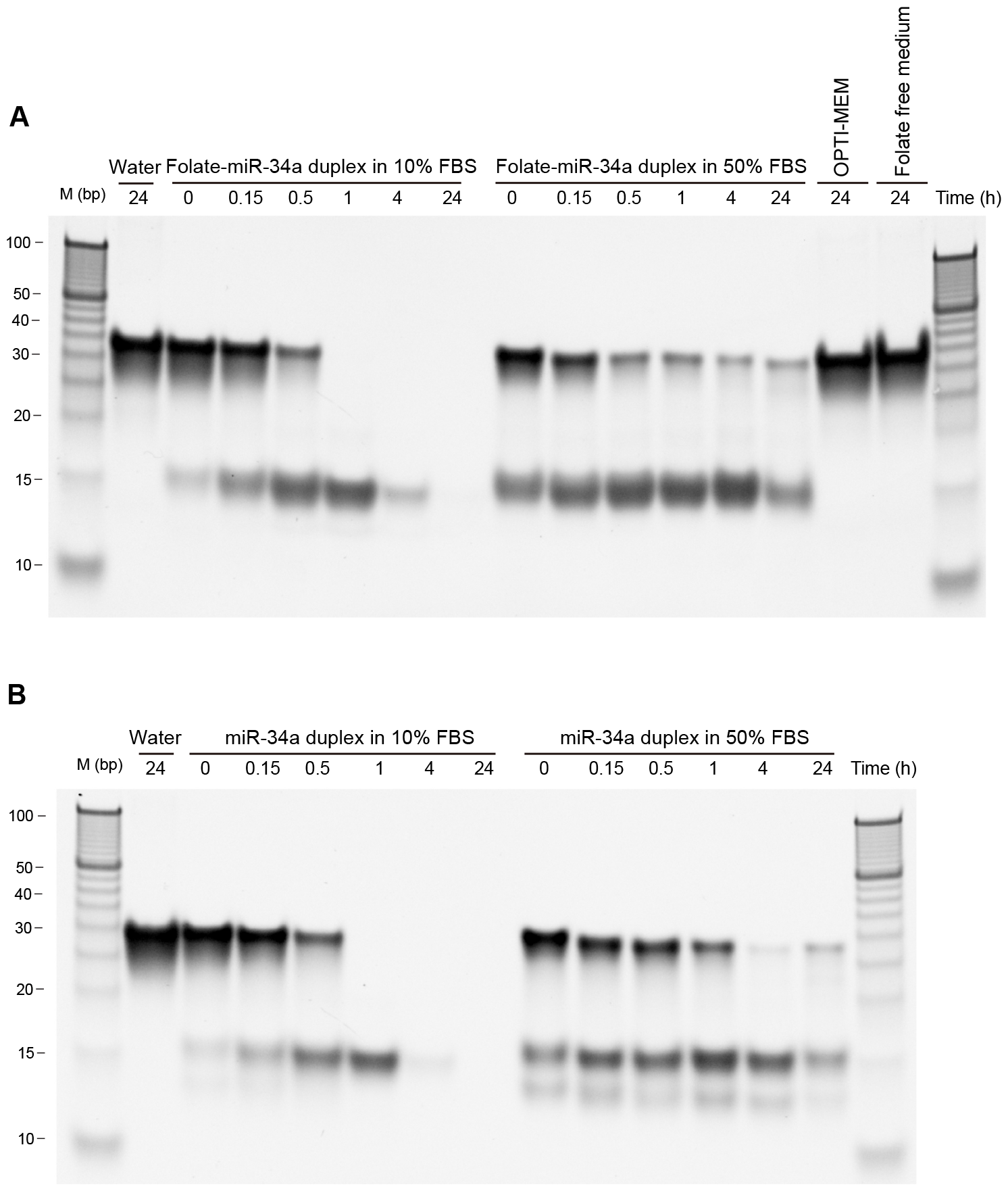

